# Crystallographic Fragment Screen of Coxsackievirus A16 2A Protease identifies new opportunities for the development of broad-spectrum anti-enterovirals

**DOI:** 10.1101/2024.04.29.591684

**Authors:** Ryan M. Lithgo, Charles W.E. Tomlinson, Michael Fairhead, Max Winokan, Warren Thompson, Conor Wild, Jasmin Cara Aschenbrenner, Blake H. Balcomb, Peter G. Marples, Anu V Chandran, Mathew Golding, Lizbe Koekemoer, Eleanor P. Williams, SiYl Wang, Xiaomin Ni, Elizabeth MacLean, Charline Giroud, Andre Schutzer Godoy, Mary-Ann Xavier, Martin Walsh, Daren Fearon, Frank von Delft

## Abstract

*Enteroviruses* are the causative agents of paediatric hand-foot-and-mouth disease, and a target for pandemic preparedness due to the risk of higher order complications in a large-scale outbreak. The 2A protease of these viruses is responsible for the self-cleavage of the poly protein, allowing for correct folding and assembly of capsid proteins in the final stages of viral replication. These 2A proteases are highly conserved between *Enterovirus* species, such as *Enterovirus A71 and Coxsackievirus A16*. Inhibition of the 2A protease deranges capsid folding and assembly, preventing formation of mature virions in host cells and making the protease a valuable target for antiviral activity. Herein, we describe a crystallographic fragment screening campaign that identified 75 fragments which bind to the 2A protease including 38 unique compounds shown to bind within the active site. These fragments reveal a path for the development of non-peptidomimetic inhibitors of the 2A protease with broad-spectrum anti-enteroviral activity.

## Introduction

*Picornaviridae Enterovirus A71* and *Coxsackie A16* are part of the group of non-polio *Enteroviruses* from the *Picornaviridae* family, they are one of the primary causative agents of hand-foot-and-mouth disease (HFMD). HFMD is usually a mild and self-limiting paediatric infection typically occurring in children under five, but can on occasion break out amongst older children and adults. Typical symptoms include fever, skin rash on the hands and feet, and mouth sores that usually abate within ten days. HFMD is fairly contagious, with droplet transmission and consumption of infected substances being common vectors for infection. In serious cases complications include viral meningitis, as well as encephalitis and polio-like paralysis in rare cases.^4,5,6,7^ These enteroviruses pose a future pandemic risk based on these factors, a small increase in infectivity, serious morbidity or adult infection based on viral evolution within reservoirs and populations could create an overwhelming amount of cases that spread rapidly amongst populations and overwhelm care resources.

*Enteroviral* 2A proteases (2A^pro^) are chymotrypsin-like cysteine proteases responsible for self-cleavage from the viral polypeptide, engendering its own correct folding and that of the capsid proteins that proceed it. This cleavage is vital in ensuring viral maturation and the progression of the viral life cycle and thus infection. 2A^PRO^ carries out secondary proteolytic interactions with host cell proteins including eukaryotic initiation factor 4 gamma (eIF4G), reducing expression of host proteins whilst viral proteins are translated unopposed through internal ribosome entry sites.^9,10,11^

*Enteroviral* 2A^pro^ and homologous enzymes from *rhinoviruses* are not widely explored as drug targets in existing art. Inhibitors that do exist are either peptidomimetics or have other undesirable characteristics for clinical candidates.^13,14 15^ Telaprevir, an FDA approved for treatment of hepatitis-C infection, binds EV-A71 2A^PRO^ and EVD68 2A^PRO^ with micromolar affinities, and has proved useful as a tool compound for investigation of both proteins however is not used as a treatment against *Enterovirus’s*.^16^

The crystallographic fragment screening approach builds upon widely used fragment screening techniques with the added benefit of three dimensional structure and interaction information and high sensitivity - essential for identifying weak, but efficient, binders that can be rapidly transformed into potent inhibitors.^19,20,23^ This structure-based method creates a rich dataset of ligand coverage information, and can lead to the discovery of potential allosteric sites and cryptic pockets that may not be identified by other biophysical screening methods.^20,21,22,23^ By using large and diverse fragment libraries, and through technological innovation in crystal soaking, harvesting, automated data collection and data processing, the fragment soaking experiment is a well proven method for delivering high quality, low affinity binders at a rapid pace.^24,25^ Discussed below are the results of a high throughput fragment screening campaign, the target enablement work required to make it possible, and the potential applications of this data in the efficient development of lead-like molecules targeting high pandemic risk viruses, such as Enterovirus A71.

## Materials & Methods

### 2A^PRO^ Protease Construct Design

A *E*.*coli* codon optimised synthetic gene for 2A was based on accessible deposited sequences from *Coxsackievirus A16* (G-10) 2A protease and ordered from TWIST Bioscience. The gene corresponds to residues 869-1012 of the polyprotein (Uniprot id:Q65900). The synthetic gene was cloned as a SUMO fusion using the golden gate method (DOI: 10.1101/2024.02.13.579886).

~~~
**MHHHHHHGSGDQEAKPSTEDLGDKKEGEYIKLKVIGQDSSEIHFKVKMTTHLKKLKESYCQ
RQGVPMNSLRFLFEGQRIADNHTPKELGMEEEDVIEVYQEQTGG**\*SGAIYVGNYRVVNRHL
ATHNDWANLVWEDSSRDLLVSSTTAQGCDTIARCDCQTGVYYCSSRRKHYPVSFSKPSLIF
VEASEYYPARYQSHLMLAVGHSEPGDCGGILRCQHGVVGIVSTGGNGLVGFADVRDLLWLD
EEAMEQ
~~~

*Sequence 1: Final construct sequence for CVA16 2A*^*PRO*,^ *The SUMO tag is displayed in bold and the CVA16 2Apro is underlined. The SUMO cleavage site is indicated by an asterisk*.

### 2A^PRO^ Protease Protein Expression and Purification

2A^PRO^ expression entailed the transformation of 100 ng of plasmid into the *E. coli* protein expression strain BL21(DE3) and a single colony used to inoculate 10 mL of 2xLB media (50 μg/mL kanamycin). The starter culture was grown overnight at 37°C with 250 rpm shaking and the following day 10 mL were used to inoculate 1 L of AIM-TB (ForMedium) in a 2.5 L baffled shake flask supplemented with 50 μg/mL kanamycin. Cells were grown for 4 h at 37°C and then for 20 h at 18°C using 250 rpm shaking. Cells were harvested by centrifugation (4000 g) and the pellet frozen at −80°C for storage.

The cell pellet was resuspended in base buffer (50mM HEPES (pH 7.5), 300 mM NaCl, 5% glycerol, 0.5mM TCEP) supplemented with 0.5 mg/mL lysozyme, 0.05 mg/mL benzonase, 2 mM MgCl, 25 mM imidazole and 2 complete™ EDTA-free Protease Inhibitor Cocktail tablets), then sonicated over ice at 40% amplitude, 4s on 12s off, for 7 minutes. 2A^pro^ was first purified by IMAC using Ni-Sepharose 6 FF resin (Cytiva) with the lysate and resin incubated for 30 mins. The resin was then washed with base buffer supplemented with 30 mM imidazole, then with 50 mM imidazole. The target protein was eluted with base buffer containing 500 mM imidazole. Eluted protein was buffer-exchanged into base buffer using HiPrep 26/10 desalting column (Cytiva). His-Ulp1 was added at 1:100 ratio to the eluate protein content to cleave the His-SUMO purification tag by incubation overnight. Reverse-IMAC was carried out the next day to purify the cleaved protein. Finally, cleaved protein was polished by size exclusion chromatography (SEC) (HiLoad 16/600 Superdex 75 pg (Cytiva)) in SEC buffer (10 mM HEPES pH 7.4, 300 mM NaCl, 5% glycerol, 0.5mM TCEP). Efficacy of the process and final protein purity was assessed by SDS-PAGE using NuPAGE 4-12 % Bis-Tris Midi Protein Gels (ThermoFisher Scientific) and mass spectrometry. Final yield of pure protein was 3 mg per litre of culture.

The whole protocol can be found here: dx.doi.org/10.17504/protocols.io.rm7vzj3y5lx1/v1.

### 2A^pro^ Fluorescence Resonance Energy Transfer (FRET) Assay

The proteolytic activity of purified CVA16 2A^pro^ was tested by determining the IC_50_ value for the control compound Telaprevir (Pubchem CID 3010818) in a biochemical FRET assay. Telaprevir was dispensed onto 384-well, black plate (Greiner, #781209) using the Echo liquid dispenser (Beckman Coulter) at a final top concentration of 200 μM using a dilution factor of 2. The IC_50_ curve includes duplicates of each concentration and solvent control. Fifty microliters of 2x (5 μM) CVA 2A^pro^ solution (50 mM Tris pH 7.0, 150 mM NaCl, 10% glycerol and 1 mM TCEP) was added to dispensed Telaprevir and incubated for 1 hour at room temperature. The reaction was initiated by the addition of 50 μL of 2x (20 μM) substrate solution (Dabcyl-TAITTLGKFGQE-Edans (LifeTein, USA)) made in the reaction buffer. The fluorescence intensity at 460 nm was read every 30 seconds for 2 hours in kinetic mode, which included a shaking step of the plate between each measurement. The IC_50_ was calculated by plotting the initial velocity against various concentrations of the tested inhibitor by using a four parameter dose−response curve in Prism (v8.0) software. The Z’ or Z-factor was calculated as below (μ: mean; σ: standard deviation, s: no inhibitor signal, highest inhibition signal).

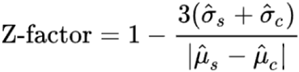

The full protocol can be found here: dx.doi.org/10.17504/protocols.io.q26g7186qgwz/v1

### 2A^PRO^ Crystallisation Condition Screening and Optimization

2A^PRO^ protein was screened for crystallisation conditions using the Shotgun (SG1) (Molecular Dimensions) and Index (Hampton Research) sparse-matrix screens at a protein concentration of 12.5 mg/mL and 18 mg/mL. Screens were set up in SwissSci 3 drop plates, with drops set up at 3 ratios 1:2, 1:1 & 2:1 (protein:reservoir) to a total volume of 300 nL. Once prepared screens were incubated at 20°C with images taken after 24 hours and then following a fibonacci sequence of timings.

A hit condition from Shotgun (SG1) that generated a single diffracting crystal was optimised using a Scorpion liquid handler (Art Robbins Instruments) to create a 96 condition screen consisting of 0.1 M MES pH 6.7, with PEG 20,000 varied between 13.5% and 18.9% w/v. The resulting screen was used as a reservoir solution (35 μL) in Swiss-Ci 3-lens midi plates, and combined with 2A^PRO^ protein in 150:150, 100:200, and 200:100 nL ratios. The majority of these conditions resulted in diffracting crystals. Seeding was utilised in further rounds of crystallisation to control the number and size of crystals following a standard procedure DOI: 10.17504/protocols.io.kxygx3nwog8j/v1

Bulk crystallisation for fragment screening was carried out using a SPT Labtech Mosquito, in 3-lens Swiss-CI midi plates containing 35 μL Reservoir Solution (0.1 M MES pH 6.7, 13.5% w/v PEG 20,000). Recombinant 2A^PRO^ (18 mg/mL) was combined with Reservoir Solution and crystal seeds (1:1000 dilution) in a 150:150:50 nL ratio. Plates were incubated in a Formulatrix Rock Imager (20 °C) with periodic imaging. Crystal formation was observed in 75% of drops within 24 hours.

### XChem Fragment Screening

Library fragments from DSiP, DSiP EUbOpen Expansion Library, Fraglites, Peplites, MiniFrags, York3D and SpotXplorer^3(35-41)^ were dispensed acoustically using Echo liquid handling (Beckman Coulter) at 25% v/v and incubated at 20 °C for 3 hours.^27^ Crystals were cryo protected with 25% glycerol and harvested and cryo-cooled using a CrystalShifter stage (Oxford Lab Technologies).^28^ Diffraction data were collected at the Diamond Light Source beamlines I04-1 and I03 using Unattended Data Collection ‘Ligand recipe’ parameters (360° rotation, 0.1°/img, 0.002 s/img and 100% transmission or 10% transmission for I04-1 and I03 respectively.

### Data Processing

Diffraction data were automatically processed using Diamond Light Source’s automated analysis pipelines and further analysis was performed through XChemExplorer.^29, 30, 31^ Initial map calculation was carried out using DIMPLE.^32^ Ligand CIF restraints were generated using GRADE.^33^ Hit identification was performed using PanDDA2 and a standard set of parameters.^34^ Refinement and model building were carried out in REFMAC and COOT via the XChemExplorer platform.^35,36,37^

## Results & Discussion

### 2A^pro^ Construct

The 2A^pro^ sequence of EV-A71 and CVA16 differ by 5 amino acids, none of which are near or predicted to affect protease activity. Therefore, with the construct designed, we are confident that any fragments obtained from the screen can be directly applied to the EV-A71 2A^pro^ and allow for further fragment elaboration. A sequence alignment of the two 2A^pro^ from EV-A71 and CVA16 highlights the sequence conservation between the two (Figure 1).

**Figure 1:**
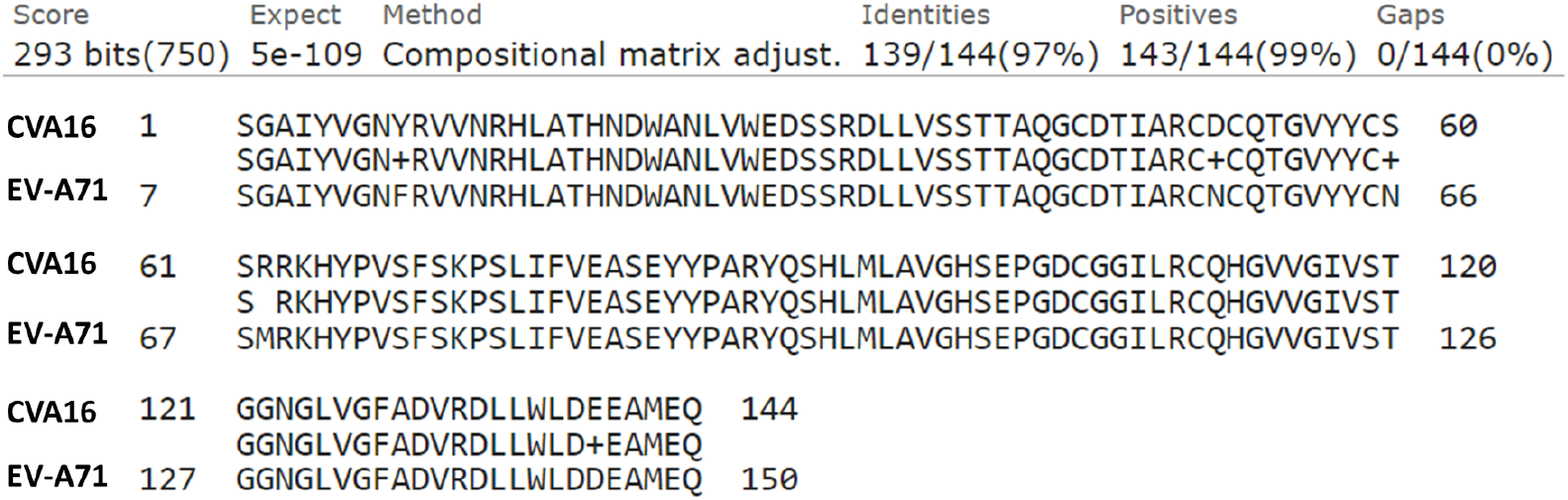
Sequence alignment of EV-A71 2A^pro^ and the 2A^pro^ sequence we designed based on the CVA16 2A^pro^ sequence.

### 2A^pro^ FRET Assay

The catalytic activity of purified 2A^pro^ was confirmed by FRET assay by measuring the IC_50_ of Telaprevir, a viral 2A protease inhibitor.^16^ The initial velocity measured in the linear range of the reaction was normalised and plotted against Telaprevir concentration. The assay shows a robust signal characterised by a signal-to background of 253, a Z’ of 0.959 and IC_50_ of 2.134 μM in the expected micromolar range (**Fig. 2**). The enzymatic activity of the protein gives confidence that the construct used for crystallographic fragment screening is biologically relevant.

**Figure 2:**
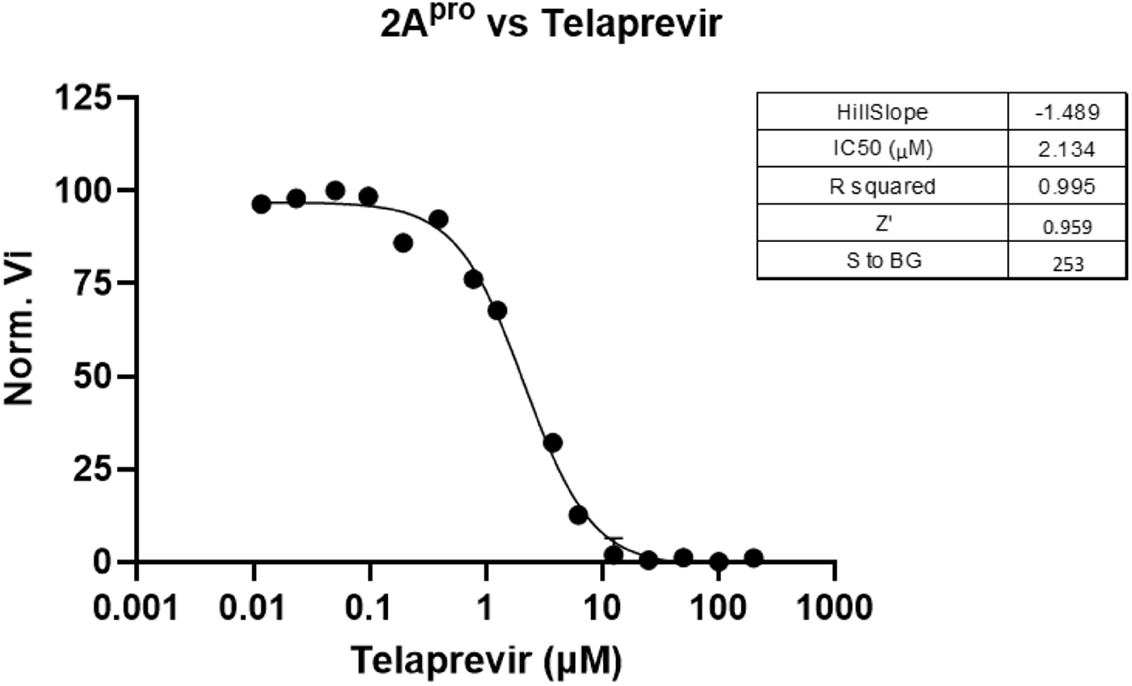
Protease assay was developed as a proof of activity of CVA16 2A^pro^ and demonstrates IC50 measurement of known inhibitor Telaprevir.

### Crystallisation of apo 2A^pro^

The crystal structure of apo 2A^pro^ was first determined in the C2 space group (a: 86.128 Å b: 57.199 Å c: 64.631 Å α: 90° β: 95.23° γ: 90°) at 1.6 Å resolution (PDB - 8POA). The asymmetric unit contains two monomeric proteins with non-crystallographic symmetry, each with a zinc ion bound as part of a structural zinc-finger motif.

The region of interest within the 2A^pro^ is the proteolytic active site. The region broadly forms a T shaped junction, which contains the catalytic Cys110, His21 & Asp39 triad residues at the centre. The density for these residues is clearly defined in the density, and within expected distances for hydrogen bonding with neighbouring residues and solvent. The 2A^pro^ active site is divided into 6 regions (S1’-S5) based on the substrate residue located in the pocket during peptide cleavage, with the catalytic triad positioned in the S1 and S2 regions (**Fig. 3-A**). The roof of the active site is a flexible loop region containing Glu88 and Tyr89, these residues govern the pocket configuration being either “open” or “closed”^10,12^, this variation in conformation is observed in the structures we obtain. The main governing residue is Tyr89, in monomer B of the apo state the tyrosine side chain is pointing into the S1 site giving a “closed”, whilst in monomer A the tyrosine is pointing away from the active site resulting in an “open” conformation (**Fig. 3-B**). The movement of the loop is seen readily across enteroviral species^8^, based on this we progressed with crystallisation and fragment screening as it was not seen as a problem.

**Fig 3:**
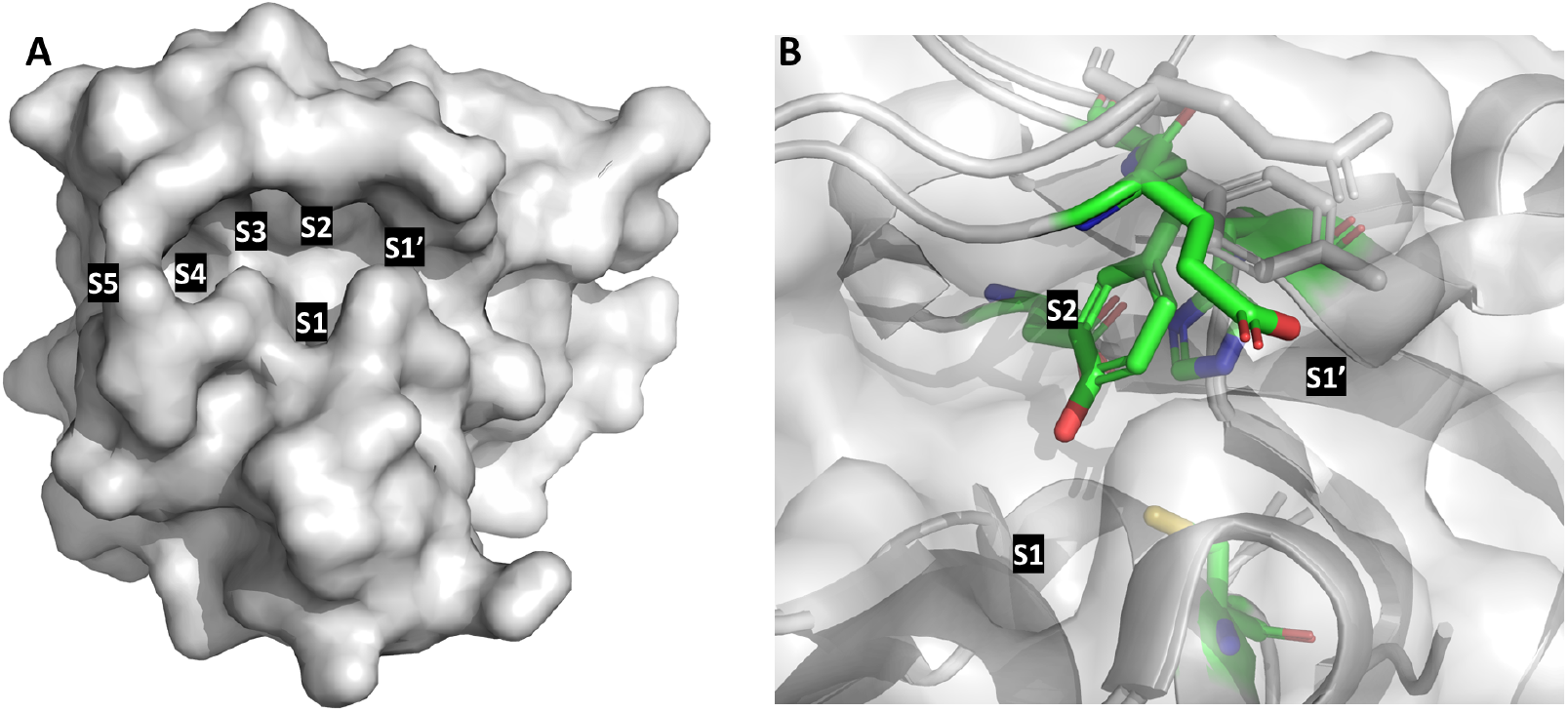
***A:*** *Top view of the “open” conformer of* 2A^pro^ showing the *active site with the subsites labelled*. ***B:*** *Overlay of the “open” (grey) and the “closed” (green) conformers of* 2A^pro^, in the closed state *Tyr89 and Glu88 can be seen pointing into the P1 subsite. Both ‘open’ and ‘closed’ forms were observed in the fragment screening data*

**Figure 3:**
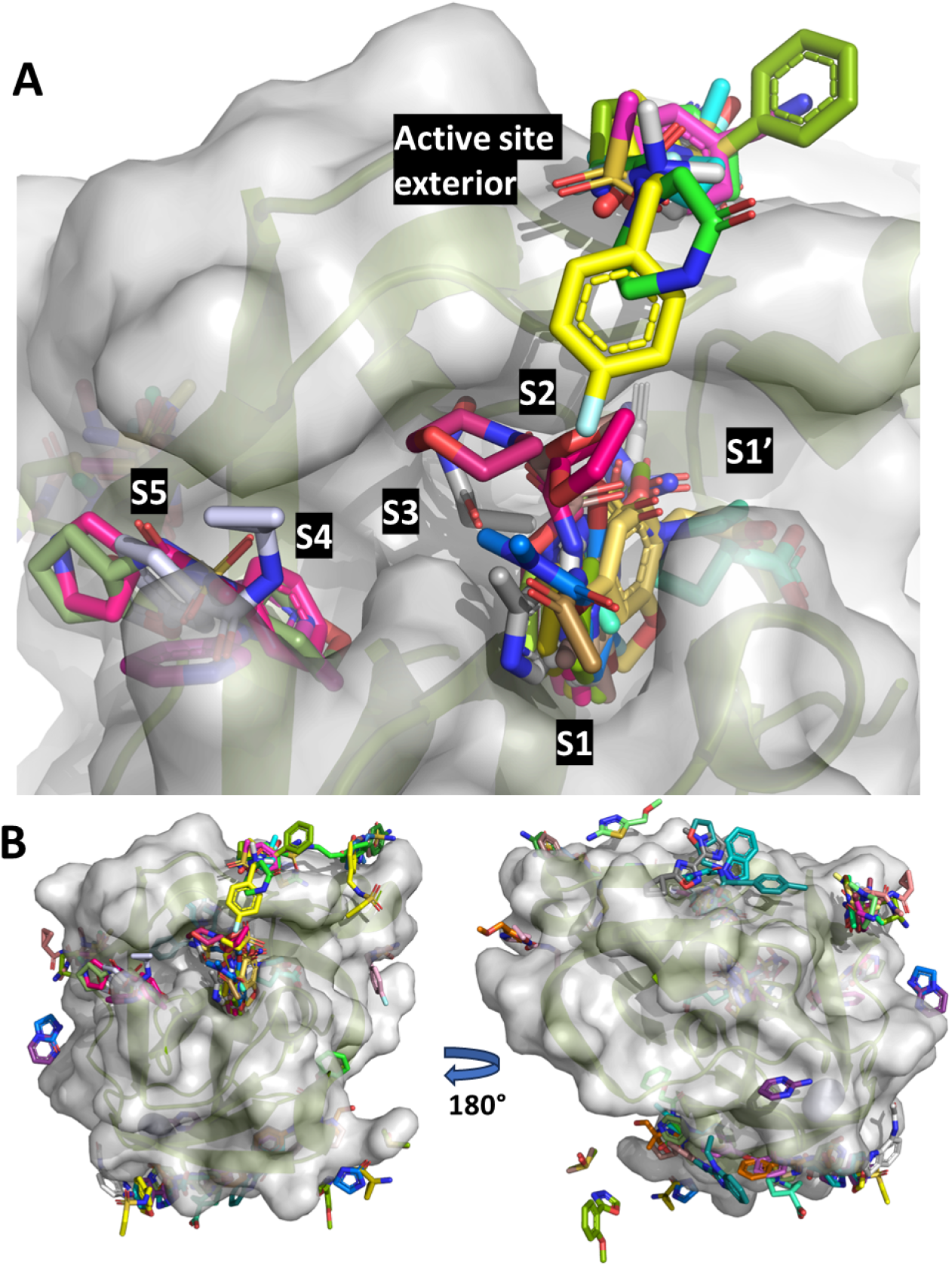
**A:** Top view of substrate 2A^PRO^ binding site with subsites labelled white-on-black moving right to left and the observed bound ligands from the fragment screen. **B:** Basal orientations showing fragment distribution across the protein surface.

Subsequent crystallisation identified an alternate C2 crystal form with a shorter cell length in the ‘C’ dimension and only a single monomer in the asymmetric unit. Since this crystal form displayed better handling characteristics and was more readily accessible for screening, it was used to produce the crystals for the fragment screen. The apo protein in this crystal form had the “closed” conformation of the active site. Throughout the screening experiment further unit cell heterogeneity was observed, with minor elongation or shortening of axes in approximately 10% of cases. It appears that these changes were within solvent channels and interfaces, as structure-to-structure the protein coordinates remained homogeneous and each was easily solved by simple molecular replacement/refinement with DIMPLE.

### Fragment Screening of 2A^pro^

Crystals were resilient to soaking with DMSO at concentrations up to 30% v/v for up to 3 hours and data quality was substantially improved by the addition of 25% v/v glycerol as a cryo-protectant. Based on a solvent concentration of 25% DMSO with 3 hour soaking, a screen of 1,129 unique fragments from DSiPoised^35^, DSiPoised EUbOpen Expansion Library^36^, Fraglites^37^, Peplites^38^, MiniFrags^39^, York3D^40^, SpotXplorer^41^ and Cov MiniFrags^46,47,48^ libraries was carried out. 891 crystals survived the soaking process, and 754 processable datasets were collected. Analysis with PanDDA2^45^ identified 2500 events, that is density which may suggest a compound is bound, however of these 102 binding events (75 unique fragments) were confidently placed across five regions (Figure 3B). The active site pocket contains 38 of these fragments, concentrated primarily in the S1 and S2 subsites (Figure 3A). A large number of fragments bind to the exterior of the active site roof, and offer potential for ligand design bridging out of the core pocket areas. Beyond that, fragments occupy four broad areas away from the active site, named Tyr64, Cys116, Trp33 and His97 based on proximity to those residues. It is possible that these regions constitute cryptic binding sites for other interactions of 2A^PRO^, though these interactions have not to date been structurally characterised.

The S1 site incorporates the 21 of the fragments from the active site events, many of which interact with Cys110 (catalytic cysteine), Ser105, Pro107, Ser125 or Gly127 through side chain or backbone interactions (**Fig. 4**). These interactions follow the broad pattern of the natural substrate (LEU) with an uncharged pocket closer to Cys110, and a charged region above for polar interactions.^12^ There are 8 additional fragments that lead into the glycine (S1 - neutral) or threonine (S2 - charged) pockets. The S2 charged pocket is created by Gln95, Tyr90, Arg93 and Asp39 sidechain interactions, and nine fragments were found to bind in this site.

**Figure 4:**
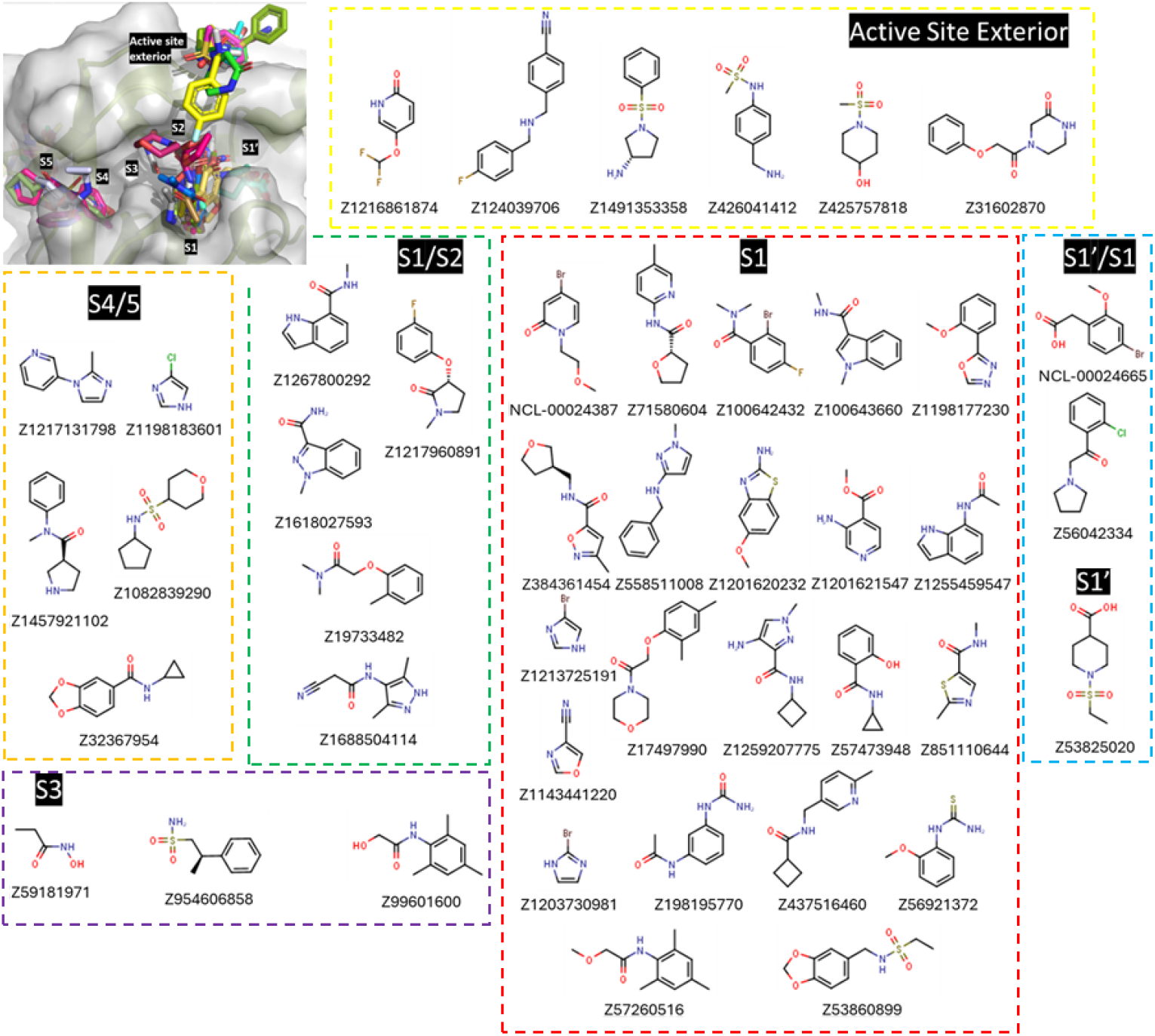
Plan view of the 2A^PRO^ S1’-S5 binding site with bound fragment chemical structures and associated ZINC IDs, with their positions within the active site.

Only 3 compounds sit wholly in the narrow S3 site, but three protrude in that direction from S2. Six compounds can be found in the broad S4/5 pocket, which opens out to the surface of the protein where interactions are primarily steric or backbone driven. Overall the 38 fragments within the active site indicates that linked or grown ligands may be able to bridge the S1’-S3 region to create strong inhibitors. A further tranche of six fragments are found above the apical loop. These so-called “active-site exterior” hits interact through the backbone atoms of residues between Ala86 and Tyr90 and may offer an additional route for expansion of elaborated molecules (**Fig. 4**).

### Structural Variation displayed by 2A^PRO^

It is clear from these data that 2A^PRO^ is fairly structurally homogeneous, and that the primary source of variation is the apical loop that controls binding site conformation. The extent of this variation is seen in Figure 5, where 107 different loop geometries are overlaid after alignment by lowest C-α RMSD. It is likely that controlling this loop geometry will be invaluable in designing inhibitor molecules, especially if the fragments binding to its top surface are incorporated into the design scheme.

**Figure 5:**
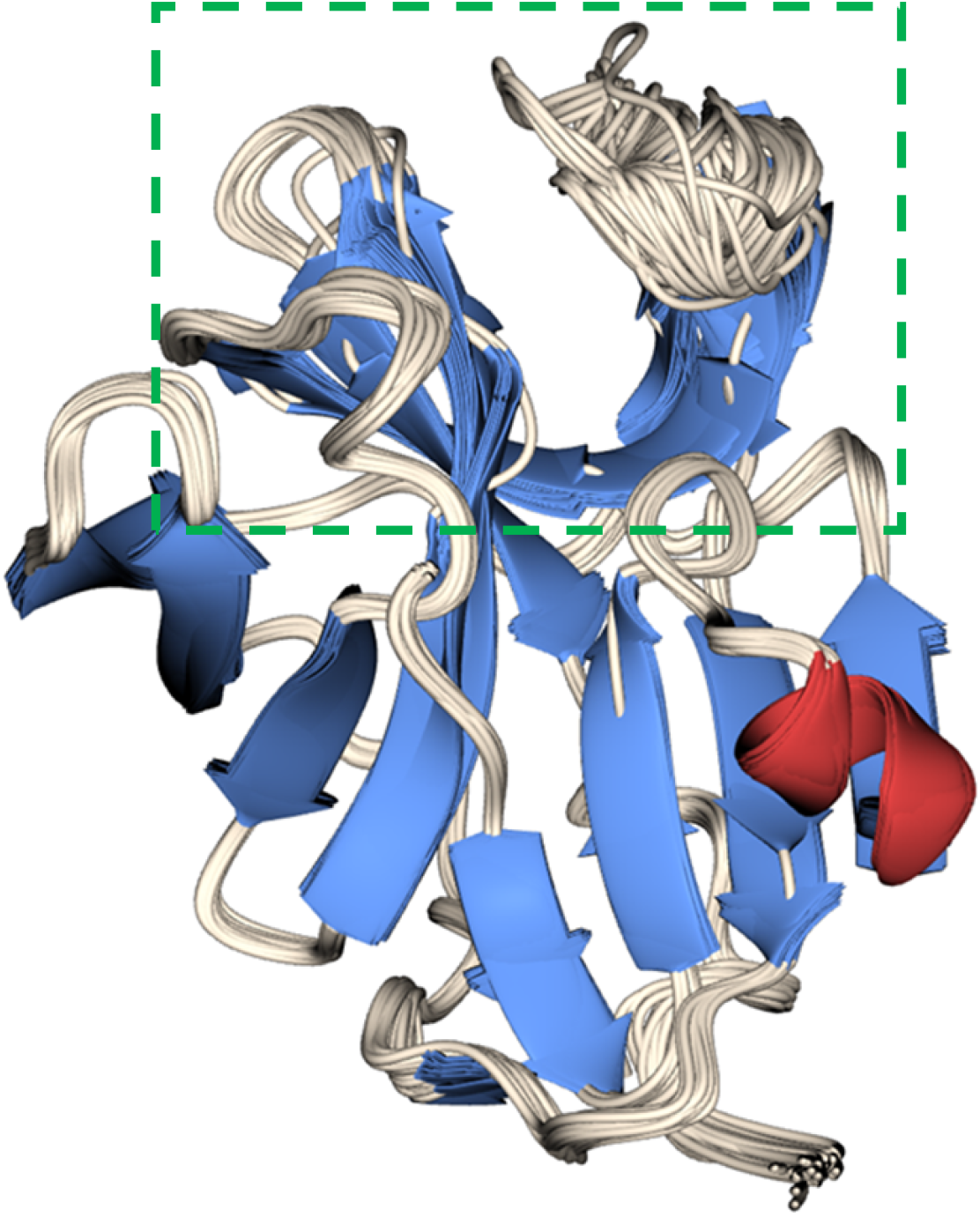
Ensemble of 107 protein chains from 2A^PRO^ showing secondary structure (β-sheet - Blue, a-helix - Red, loops and unstructured - white). The active site channel can be seen at the top of the structure (Green Box) with S1’ closest to the viewer. The apical loop X-X which governs active site conformation displays the greatest structural variation between structures.

## Conclusions

By utilising a readily accessible 2A^PRO^ construct that illustrates the high sequence similarity of the 2A^PRO^ between enteroviral species, we were able to rapidly perform a crystallographic fragment screen that promotes confidence in the potential design of pan-enteroviral inhibitors.

Initial crystallisation screening of 2A^PRO^ readily supplied well diffracting crystals in conditions that were efficiently optimised using simple buffer components. Additional seeding helped to control the crystallisation and provide bulk crystals ready for high throughput crystallographic fragment screening. The fragment screen has afforded an excellent set of 75 discrete fragments covering six key areas of the protein. The 38 fragments within the active site engage in interactions with key substrate binding residues, as well as spanning most of the S1’-S5 subsites involved in natural substrate binding. The large number of crystallographic datasets also provides insight into the conformational variation of the protein, which appears to be limited to the apical loop that controls binding site conformation. Understanding this conformational shift will improve the likelihood of positive outcomes in fragment-hit-lead development cycles ahead. Structures of fragment bound 2A^PRO^ were made publicly available on the PDB in January 2024, and are also available to view in Fragalysis which offers a user-friendly interface for viewing and disseminating the aligned structures of each fragment bound structure.

It is highly likely that fragment-merging and elaboration can be utilised to efficiently develop compounds that span the entirety of the active site whilst keeping true to the parent hit structures. These future compounds would hopefully achieve high binding potencies and would suitably inhibit the natural function of the protein *in vivo*.

## Supporting information

## Acknowledgements

The authors would like to acknowledge Diamond Light Source for access to the fragment screening facility XChem, for usage of DSi-Poised and other libraries and for beamtime on the I04-1 beamline. We would also like to thank Neil Paterson and Mark Williams for their assistance with data collection of the I03 beamline.

## Author contributions

RML and CWET carried out fragment screening, data analysis and writing the manuscript. DF, JCA, BHB, and PGM supported data processing, data collection and validation. MW and WT assisted with computational analysis and chemistry. SW, EW, MF, XN, CG and LK carried out protein expression, purification and quality control. DF, LK and ASG reviewed the manuscript.

## Data and materials availability

All data needed to evaluate the conclusions in the paper are present in the paper and/or the Supplementary Materials. Table 1 for crystallographic stats is provided as an .csv file.

Structures of CVA16 2^PRO^ were deposited to the PDB under accession codes: 7H2T, 7H2U, 7H2V, 7H2W, 7H2X, 7H2Y, 7H2Z, 7H30, 7H31, 7H32, 7H33, 7H34, 7H35, 7H36, 7H37, 7H38, 7H39, 7H3A, 7H3B, 7H3C, 7H3D, 7H3E, 7H3F, 7H3G, 7H3H, 7H3I, 7H3J, 7H3K, 7H3L, 7H3M, 7H3N, 7H3O, 7H3P, 7H3Q, 7H3R, 7H3S, 7H3T, 7H3U, 7H3V, 7H3W, 7H3X, 7H3Y, 7H3Z, 7H40, 7H41, 7H42, 7H43, 7H44, 7H45, 7H46, 7H47, 7H48, 7H49, 7H4A, 7H4B, 7H4C, 7H4D, 7H4E, 7H4F, 7H4G, 7H4H, 7H4I, 7H4J, 7H4K, 7H4L, 7H4M, 7H4N, 7H4O, 7H4P, 7H4Q, 7H4R, 7H4S, 7H4T, 7H4U, 7H4V, 7H4W, 7H4X, 7H4Y, 7H4Z, 7H50, 7H51, 7H52, 7H53, 7H54, 7H55

Aligned ligand structures are also available through the Fragalysis platform for visualisation and analysis.

